# Patterns of brain asymmetry associated with polygenic risks for autism and schizophrenia implicate language and executive functions but not brain masculinization

**DOI:** 10.1101/2021.03.19.436120

**Authors:** Zhiqiang Sha, Dick Schijven, Clyde Francks

**Affiliations:** Language and Genetics Department, Max Planck Institute for Psycholinguistics, Nijmegen, The Netherlands; Donders Institute for Brain, Cognition and Behaviour, Radboud University, Nijmegen, The Netherlands

## Abstract

Autism spectrum disorder (ASD) and schizophrenia have been conceived as partly opposing disorders in terms of systemizing versus empathizing cognitive styles, with resemblances to male versus female average sex differences. Left-right asymmetry of the brain is an important aspect of its organization that shows average differences between the sexes, and can be altered in both ASD and schizophrenia. Here we mapped multivariate associations of polygenic risk scores (PRS) for ASD and schizophrenia with asymmetries of regional cerebral cortical surface area, thickness and subcortical volume measures in 32,256 participants from the UK Biobank. PRS for the two disorders were positively correlated (r=0.08, p=7.13×10^−50^), and both were higher in females compared to males, consistent with biased participation against higher-risk males. Each PRS was associated with multivariate brain asymmetry after adjusting for sex, ASD PRS r=0.03, p=2.17×10^−9^, schizophrenia PRS r=0.04, p=2.61×10^−11^, but the multivariate patterns were mostly distinct for the two PRS, and neither resembled average sex differences. Annotation based on meta-analyzed functional imaging data showed that both PRS were associated with asymmetries of regions important for language and executive functions, consistent with behavioural associations that arose in phenome-wide association analysis. Overall, the results indicate that distinct patterns of subtly altered brain asymmetry may be functionally relevant manifestations of polygenic risk for ASD and schizophrenia, but do not support brain masculinization or feminization in their etiologies.

## Introduction

ASD is a childhood-onset disorder that features deficits in social communication and social interaction, together with restricted or repetitive behaviour^1^. The cognitive and behavioural profile of ASD has been conceived in terms of over-systemizing (focusing heavily on the component variables of systems) and under-empathizing (a deficit in recognizing the thoughts and emotions of others)^2^. A similar concept involves over-mechanizing in ASD (a focus on interaction with the physical environment) and under-mentalizing (deficits in attributing mental states to others)^3^. Given that average sex differences also exist in these dimensions, with males tending to outperform on systemizing tasks and underperform on empathizing tasks, ASD was proposed to resemble excessive masculinization of the brain^2^.

In contrast, positive symptoms in schizophrenia, which can involve the incorrect attribution of thoughts or feelings to external agents, have been conceived in terms of hyper-mentalizing^3–6^. This has led to models of psychosis and ASD as opposing disorders of the social brain that partly mirror average sex differences^3–6^.

There are also features often found in common between ASD and schizophrenia, including impaired executive functions^7–10^ and deficits in social cognition and competence in the verbal and non-verbal domains^11, 12^. However, such similarities may be relatively superficial and not reflect etiological commonalities^11,13,14^. Large-scale cross-disorder analyses have pointed to shared genetic contributions to ASD and schizophrenia^15^, but while these have been particularly evident for rare mutations of relatively high penetrance^16,17^, the genetic correlation based on common single nucleotide polymorphisms (SNPs) has been estimated at only roughly 0.2 or less^18^. Therefore, both shared and independent genetic contributions are involved.

Left-right asymmetry is a pervasive organizing principle of human brain structure and function^19,20^, including for networks involved in language and social cognition^21–24^, executive^25^ and affective processes^26,27^. Various aspects of hemispheric asymmetry can be altered in ASD or schizophrenia, which include regional anatomical measures of grey and white matter, structural and functional connectivity, and behavioural associations (left-handedness has an elevated frequency in both disorders compared to the general population)^28–42^. Some structural and functional measures of hemispheric asymmetry also show average differences between the sexes^43–45^, and foetal testosterone levels relate to the development of grey matter asymmetries of some cerebral cortical regions^46^. Population-average hemispheric asymmetries are established prenatally^47,48^, likely through a genetically defined program^49–51^, and variation in some aspects of brain structural asymmetry is significantly heritable^43,44^. Therefore, brain asymmetry presents a potential intermediate phenotype between genes and diagnosis, which can help to identify etiologic similarities and differences between ASD and schizophrenia, including with respect to masculinization versus feminization.

In 32,256 adult participants from the general population UK Biobank dataset, we previously found that 42 regional brain asymmetry indexes (AIs) showed significant SNP-based heritabilities ranging from 2.2 to 9.4%^52^. These comprised 28 cortical surface area AIs, 8 cortical thickness AIs, and 6 subcortical volume AIs, where the AI for a given region and individual was calculated as (Left-Right)/((Left+Right)/2)^52^. Multivariate genome-wide association scanning (GWAS) of these AIs implicated genes involved in microtubule-related functions, and genes particularly expressed in the embryonic and fetal brain^52^. These observations are compatible with known roles of the cytoskeleton in shaping cellular chirality, which can initiate left-right asymmetry in the development of other organs of other species^53–58^. SNPs that showed relatively low multivariate association p values in relation to brain asymmetries^52^ also tended to show low p values in publicly available GWAS summary statistics for ASD and schizophrenia, which indicates a genetic overlap of brain asymmetries with these disorders.

Substantial SNP-based heritabilities for both ASD and schizophrenia^18^ indicate that they often arise at the severe tail of a continuous scale of liability across the general population^59^. One way to approach the etiologies of these disorders is therefore to study associations of polygenic risk with brain structure and function in general population datasets. In this approach, the polygenic risk of a given disorder for each individual within a population cohort is estimated from their own genotypes, in combination with SNP-wise summary statistics from large-scale GWAS for that disorder^60–65^.

Here we made use of the UK Biobank imaging genetics dataset to compare and contrast the associations of PRS for ASD and schizophrenia with brain structural asymmetry in the general adult population. We then queried meta-analyzed data from functional magnetic resonance imaging (fMRI) studies, to annotate functionally the regions showing the strongest associations of their asymmetries with each disorder PRS. We also investigated whether either of the two disorder PRS is associated with a more male-like or female-like pattern of brain asymmetry. Finally we performed phenome-wide association analyses for the two disorder PRS, to understand which other brain, behavioural, clinical and other types of variables are related to polygenic risk for these disorders in this dataset.

## Methods

### Participants

The UK Biobank is a general adult population cohort^66^. Data availability and processing (below) resulted in 32,256 participants (15,288 male, 16,968 female) for the PRS-imaging analysis. The age range was 45-81 years (mean 63.77).

### Genetic quality control

We used imputed SNP genotype data released by the UK Biobank (March 2018). We excluded a random member of each pair of individuals with estimated kinship coefficient >0.442^66^, participants with a mismatch of their self-reported and genetically inferred sex, putative sex chromosome aneuploidies, principle component-corrected heterozygosity >0.19, or genotype missing rate >0.05^66^. Analyses were restricted to participants with ‘white British ancestry’ as defined by Bycroft et al.^66^. We retained 9,803,522 bi-allelic variants with minor allele frequencies >1%, INFO score ≥ 0.7, and Hardy-Weinberg equilibrium p-value ≥ 10^−7^.

### Neuroimaging

Details of image processing and quality control are described elsewhere^67^. Measures of regional cortical surface area, cortical thickness and subcortical volumes were derived from T1-weighted MRI scans (Siemens Skyra 3 Tesla MRI with 32-channel radio frequency receive head coil), and released by the UK Biobank (February 2020, full protocol: http://biobank.ndph.ox.ac.uk/showcase/refer.cgi?id=2367). Briefly, Freesurfer 6.0 was used to parcellate the cerebral cortex into 34 regions per hemisphere according to the Desikan-Killiany atlas^68^, and to segment 7 subcortical structures per hemisphere. Cortical surface area was measured at the grey-white matter boundary, and cortical thickness was measured as the average distance in a region between the white matter and pial surfaces. Data for the temporal pole were excluded as unreliable^67^. In addition, separately per measure, we removed data-points greater than six standard deviations from the mean.

We calculated the AI for each matching pair of left and right measures, in each participant, as (left-right)/((left+right)/2). We then applied rank-based inverse normalization followed by linear regression to remove shared variance with potential confound variables, i.e. age, non-linear age (age-mean_age)^2^, the first ten principle components capturing genome-wide diversity in the genotype data^66^, X-, Y- and Z-scanner position parameters, T1 signal-to-noise ratio, T1 contrast-to-noise ratio, assessment center, genotyping array, and sex. We previously found that only 42 of the AIs showed significant SNP-based heritabilities in this dataset (false discovery rate, FDR<0.05)^52^. Accordingly, only these 42 AIs were analyzed in the present study (Supplementary Table 1).

### Polygenic risk and brain asymmetry

We downloaded genome-wide, SNP-wise summary statistics from GWAS studies of ASD^69^ (n=46,350) and schizophrenia^36^ (n=82,315). Separately for the two disorders, we used PRS-CS^70^ to calculate PRS for each individual in the UK Biobank imaging dataset. PRS-CS uses a high-dimensional Bayesian regression framework to infer posterior effect sizes of SNPs, using genome-wide association summary statistics (i.e. for the 22 autosomes, excluding the sex chromosomes). We used default parameters and the recommended global effect size shrinkage parameter phi=0.01, together with linkage disequilibrium (LD) information based on the 1000 Genomes Project phase 3 European-descent reference panel^71^. The ASD PRS was based on 1,092,064 autosomal SNPs, and the schizophrenia PRS was based on 1,097,357 autosomal SNPs (these numbers came from the 3-way overlap between UK Biobank data, disorder GWAS data and 1000 Genomes data). PRS-CS has been shown to perform highly similarly to other, recently-developed PRS methods for the prediction of psychiatric diseases, and noticeably better than using LD-based clumping with P-value thresholding^72^. Both PRS were z-score scaled for subsequent analyses.

Separately for each PRS, canonical correlation analysis (CCA) (‘canoncorr’ function in MATLAB: https://nl.mathworks.com/help/stats/canoncorr.html) was used to test for its multivariate association with the 42 brain AIs. This analysis found the linear combination of AIs (i.e. the canonical AI) that was maximally associated with a given PRS across individuals.

Post hoc univariate testing was performed by Pearson correlation between each PRS and unilateral brain measures (i.e. separately for left and right measures, adjusted for the same covariate effects as above).

### Comparing multivariate patterns of brain asymmetry associated with polygenic risk for ASD and schizophrenia

The above CCA analyses resulted in loadings, one value per brain regional AI, that reflected the extent and direction to which each AI drove the multivariate association with a given disorder PRS. The loadings were computed as correlations between each AI and the canonical variable representing all AIs in a given CCA, across the 32,256 individuals.

We then calculated the correlation between the 42 loadings (one per AI) derived from the CCAs of the two disorder PRS. This correlation indicated the extent to which PRS for the two disorders were associated with similar or contrasting patterns of multivariate brain asymmetry. The empirical p value of this correlation was calculated through 10,000 permutations in which within-subject relations between the two PRS were maintained, and within-subject relations between the 42 AIs were maintained, but the PRS were randomly shuffled with respect to the AIs.

### Functional annotation of brain regions

We identified AIs that showed the strongest loadings in CCA with respect to the ASD PRS, as those AIs having loadings >0.2 or <−0.2. The bilateral brain regions corresponding to these AIs were used to define a single mask encompassing all of these regions in MNI standard space. This mask was then analyzed using the ‘decoder’ function of the Neurosynth database (http://neurosynth.org), a platform for large-scale synthesis of task fMRI data^73^. Neurosynth uses text-mining to detect frequently used terms as proxies for concepts of interest in the neuroimaging literature: terms that occur at a high frequency in a given study are associated with all activation coordinates in that publication, allowing for automated term-based meta-analysis. We queried the database in December 2020 when it included 507,891 activation peaks reported in 14,371 studies. The input mask was first used to define a brain-wide co-activation map for that mask based on all studies in the database, and the co-activation map was then correlated with each of 1307 term-specific maps^73^. This analysis does not employ inferential statistical testing, but rather is designed to assess functional terms with respect to how strongly their meta-analyzed activation patterns correlate with a particular co-activation map derived from an input mask. We report only cognitive and disorder terms with correlation coefficients r>0.2, while excluding anatomical terms, non-specific terms (e.g. ‘Tasks’), and one from each pair of virtually duplicated terms (such as ‘Words’ and ‘Word’).

This analysis was also performed for the brain regional AIs that showed the strongest loadings in CCA with respect to the schizophrenia PRS (i.e. again for those AIs having loadings >0.2 or <−0.2).

### Analysis with respect to sex

We first tested associations of the ASD PRS and schizophrenia PRS with sex (binary integer) in the 32,256 individuals of the UK Biobank dataset, using two-sample t tests.

We then used linear regression to adjust the 42 brain AIs for potential confound effects as described above, except that sex was not included this time as a confound variable. For each of the 42 adjusted AIs, we performed a t test in relation to sex.

We then tested the correlation between the 42 loadings (one per AI) from CCA of the ASD PRS and AIs with the 42 t values from the tests of association with sex. The significance of this correlation was assessed using permutation (n=10,000) in which sex was shuffled across individuals. We further tested the correlation between the 42 loadings from CCA of the schizophrenia PRS and AIs with the

42 t values from the tests of association with sex, and again significance was assessed using permutation (n=10,000). These analyses would reveal whether either disorder PRS was associated with an overall pattern of brain asymmetry resembling a more male or female pattern.

### Phenome-wide association analysis of disorder PRS

The ASD PRS and schizophrenia PRS were separately tested as independent variables in phenome-wide association analysis in the 32,256 individuals, using PHESANT^74^. The UK Biobank has collected diverse data on sociodemographic and lifestyle factors, psychological factors, imaging traits, cognitive functioning and health. A selection of 3239 phenotypes of potential relevance to imaging genetics studies were available as part of our research application 16066. Dependent variables of continuous, binary, ordered categorical, and unordered categorical types were tested using linear, logistic, ordered logistic, and multinominal logistic regression, respectively. Prior to testing, inverse normal rank transformation was applied to continuous variables. All analyses were adjusted for the same potential confounds as further above, including sex. We excluded phenotypes that were used to construct the AIs in this study, and those that were treated as covariates in the analysis. We controlled for multiple testing using a FDR threshold 0.05, separately for the two PRS.

## Results

### Multivariate associations of brain asymmetry with disorder PRS

Frequency histograms of the two PRS are in Supplementary Figure 1. There was a low but significant correlation between the ASD PRS and schizophrenia PRS in the 32,256 participants from the UK Biobank (r=0.08, p=7.13×10^−50^).

**Figure 1.**
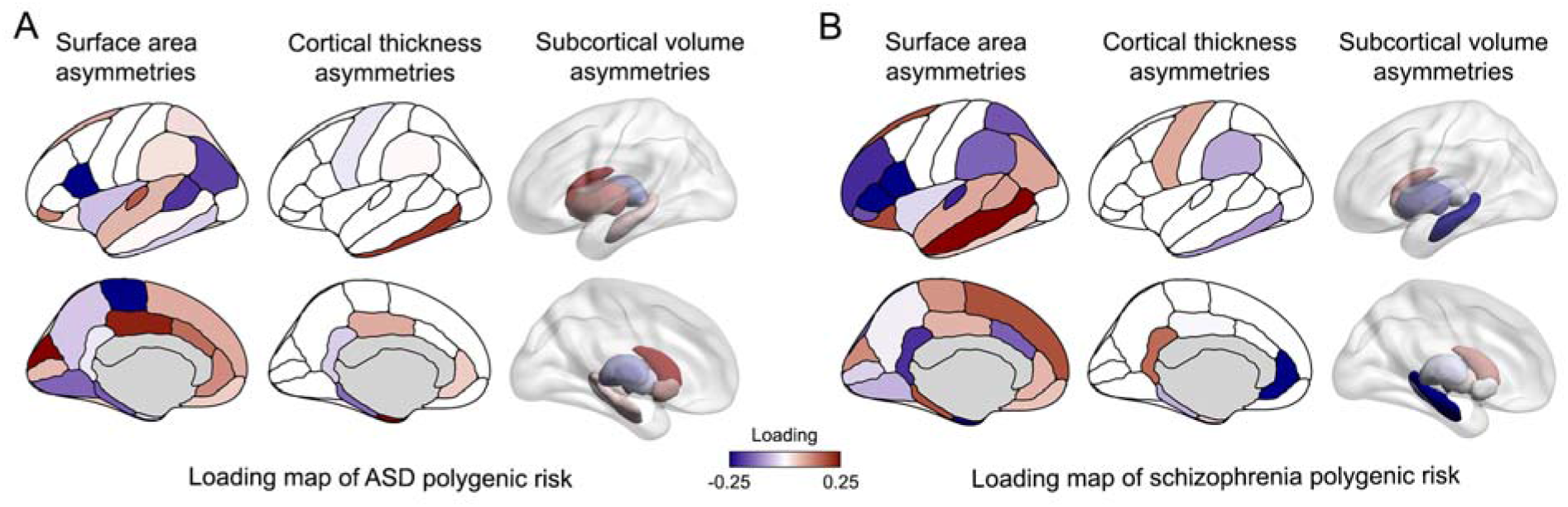
Map of associations of regional brain asymmetries with PRS for ASD and schizophrenia. Loadings of regional brain asymmetries derived from canonical correlation analyses with ASD (A) and schizophrenia (B) polygenic risk scores. A positive loading (red) for a given region indicates a leftward shift of asymmetry associated with increased polygenic risk. Conversely, a negative loading (blue) indicates a rightward shift of asymmetry associated with increased polygenic risk.

The ASD PRS showed a significant multivariate association with brain AIs, canonical correlation r=0.03, p=2.17×10^−9^. The schizophrenia PRS also showed a significant multivariate association with brain AIs, canonical correlation r=0.04, p=2.61×10^−11^. Loadings for each AI in these two CCA analyses are in Supplementary Table 2, and Figure 1A. A positive loading for a given AI indicates a leftward shift of asymmetry associated with increased polygenic risk, for a given disorder.

Conversely, a negative loading indicates a rightward shift of asymmetry associated with increased polygenic risk, for a given disorder.

There was no significant correlation (r=−0.03, empirical p=0.87) between the loadings of AIs from the CCA with ASD PRS and the CCA with schizophrenia PRS. This indicates that, considered over all regional AIs, the ASD PRS and schizophrenia PRS have generally unrelated patterns of association with brain asymmetry. Only one AI, for the surface area of the pars opercularis, showed concordant loadings with respect to both of the PRS (ASD PRS CCA loading −0.30, schizophrenia PRS CCA loading −0.28), when concordance was assessed as having both loadings >0.2, or both <−0.2 (Supplementary Table 2). No AIs with discordant loadings were observed, with loading >0.2 in one CCA and <−0.2 in the other. Results from post hoc, univariate analysis of the separate left and right brain regional measures (adjusting for the same covariates as above) are in Supplementary Table 3.

### Functional annotation of regions driving multivariate associations with the two disorder PRS

There were 8 AIs that showed loadings >0.2 or <−0.2 in CCA analysis with the ASD PRS, distributed in temporal, inferior frontal, and posterior medial cortex, as well as the caudate nucleus volume AI (Figure 2A and Supplementary Table 2). The 16 regions (8 per hemisphere) corresponding to these 8 AIs were used to create a single binary mask that was used to query the Neurosynth database of 14,371 fMRI studies (see Methods and Figure 2A). A brain-wide co-activation map (Figure 2B) was generated for this mask, based on all functional maps in the database. There were 19 term-based correlations >0.2 with the co-activation map (Figure 2C and Supplementary Table 4). The strongest of these was ‘word’ (r=0.37), and there were fourteen other terms related to language in the list, such as ‘phonological’ (r=0.36), ‘language’ (r=0.34) and ‘reading’ (r=0.33). Three terms pertaining to executive functions were also in the list: ‘demands’ (r=0.30), ‘working memory’ (r=0.24), and ‘working’ (r=0.23), as well as the term ‘ASD’ (r=0.27; Supplementary Table 4).

**Figure 2.**
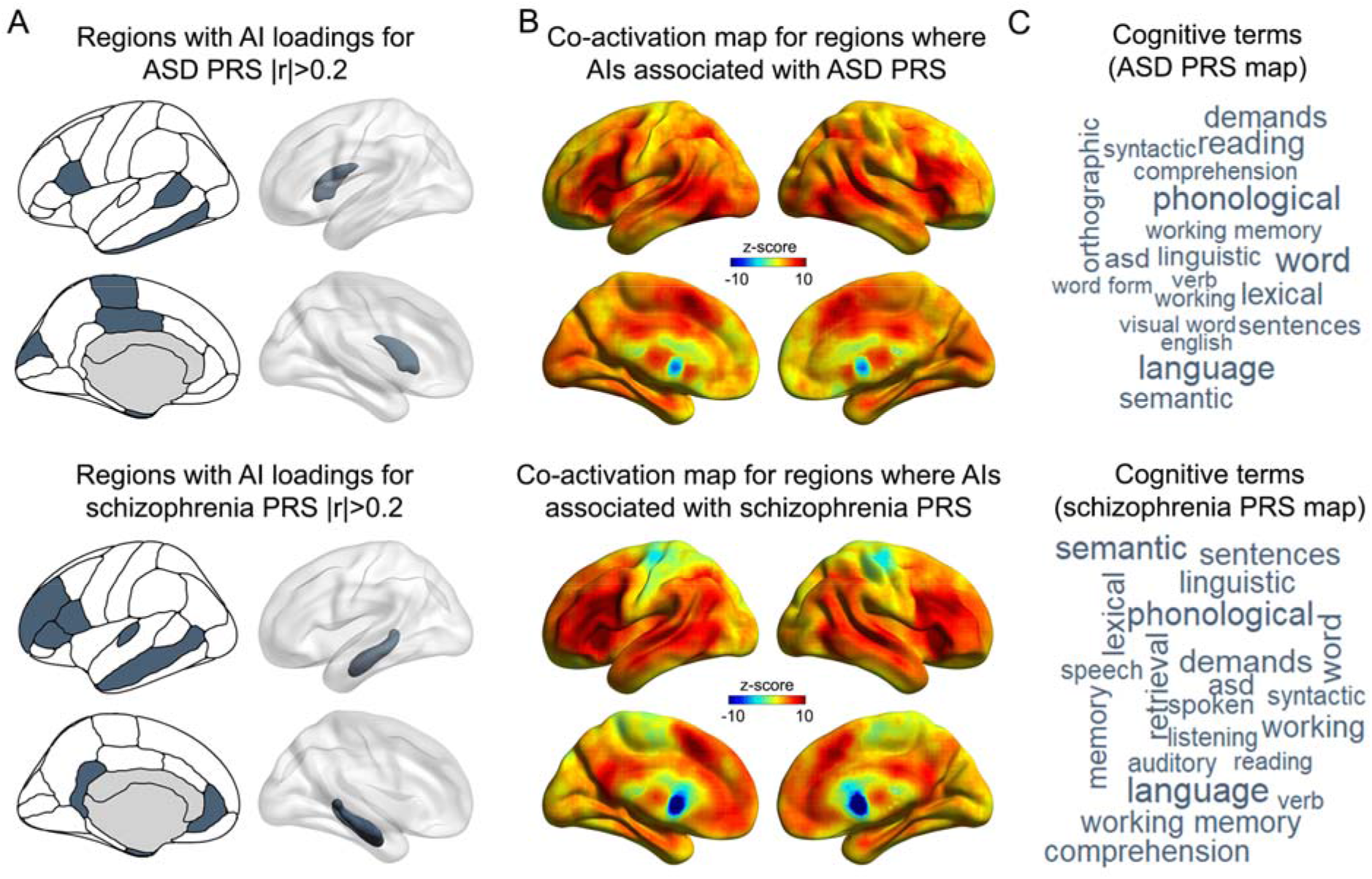
Functional annotation of regions showing the strongest associations with disorder PRS. (A) Regions for which the AIs had loadings >0.2 in canonical correlation analysis with the ASD (top) and schizophrenia (bottom) polygenic risk scores, which were used to define binary masks to query the Neurosynth database. (B) Brain co-activation maps derived from the “decoder” function of Neurosynth, corresponding to the input masks for ASD (top) and schizophrenia (bottom) polygenic risk scores. (C) Word clouds of cognitive terms associated with the co-activation maps for ASD (top) and schizophrenia (bottom) polygenic risk scores. The font sizes of the terms indicate the correlations of their corresponding meta-analytic maps with the co-activation maps.

We also created a mask corresponding to the 9 regional AIs that showed loadings >0.2 or <−0.2 in CCA analysis with the schizophrenia PRS. These were distributed over temporal, frontal and cingulate cortex, plus the hippocampus (Figure 2A). In the Neurosynth analysis there were 21 correlations greater than 0.2 (Figure 2B and 2C, Supplementary Table 4), and these were again most prominently related to language (e.g. ‘language’, r=0.33; ‘phonological’, r=0.31) and executive functions (e.g. ‘demands’, r=0.29; ‘working memory’, r=0.27), and also again included the term ‘ASD’ (r=0.26) (Supplementary Table 4), despite that most regions were different for the ASD PRS and schizophrenia PRS binary masks (Figure 2).

### Brain asymmetry, disorder PRS, and sex

Sex was associated with the ASD PRS (t=3.55, p=3.84×10^−4^) and the schizophrenia PRS (t=2.77, p=0.006), such that females tended to have higher PRS for both disorders (despite that the PRS were calculated from autosomal genotypes only). This may reflect that participation in the UK Biobank was influenced by the two PRS in a sex-dependent manner (see Discussion).

Sex was significantly associated with the majority of AIs (adjusted for all covariates as above, with the exception of sex) (Figure 3, Supplementary Table 5). In this analysis, a positive t score for a given AI indicated an average rightward shift of asymmetry in males compared to females, and vice versa. The overall map of sex associations with regional brain asymmetries (Figure 3) corresponded well with previous large-scale meta-analysis in other data ^75^.

**Figure 3.**
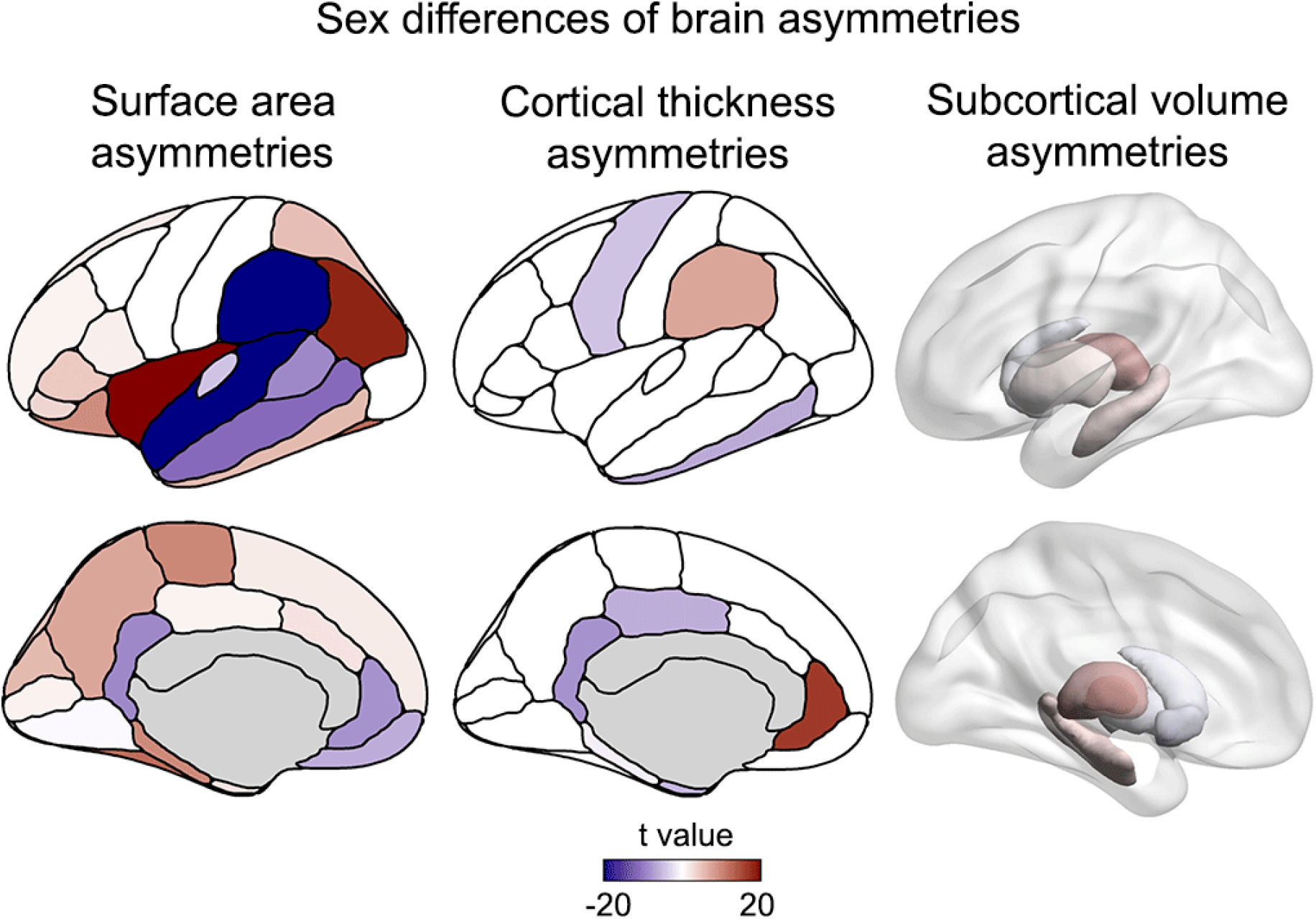
Sex differences of regional brain asymmetries. Sex differences of regional surface area asymmetries (left), cortical thickness asymmetries (middle) and subcortical volume asymmetries (right). A positive t-value (red) indicates an average leftward shift of asymmetry in females compared to males, while a negative t-value (blue) indicates an average leftward shift of asymmetry in males compared to females.

The t values from the sex-AI association analysis were not significantly correlated with loadings from the ASD PRS-AI multivariate analysis (r=−0.22, empirical p=0.16) (where sex was controlled as one of the covariates in the latter). There was therefore no evidence that ASD PRS was associated with an overall pattern of brain asymmetry more like either sex. Likewise, t values from the sex-AI analysis were not significantly correlated with loadings from the schizophrenia PRS-AI multivariate analysis (r=−0.13, empirical p=0.50; Figure 3) (where sex was again controlled as one of the covariates in the latter). This indicates that the schizophrenia PRS is also not associated with an overall pattern of brain asymmetry more like either sex.

### Phenome-wide association analyses of disorder PRS

In phenome-wide association analysis of the ASD PRS, there were 4 significant associations at FDR 0.05 (Supplementary Table 6). The most significant was a positive association with ‘Hearing difficulty/problems with background noise’ (beta=0.06, p=1.05×10^−6^), which was assessed through a touchscreen question ‘Do you find it difficult to follow a conversation if there is background noise (such as TV, radio, children playing)’. The second most significant was a positive association with ‘Townsend deprivation index’ (beta=0.03, p=1.70×10^−6^). The third was a positive association with ‘Qualifications: College or University degree’ (beta=0.05, p=5.14×10^−6^), and the fourth was a positive association with ‘Long-standing illness, disability or infirmity’ (beta=0.06, p=1.89×10^−5^).

There were 495 significant associations with the schizophrenia PRS at FDR 0.05 (Supplementary Table 7). Most significant among these were associations of higher PRS with poorer cognitive performance, e.g. ‘Interval between previous point and current one in alphanumeric path (trail #2)’ (beta=0.10, p=6.59×10^−51^), and ‘fluid intelligence score’ (beta=−0.14, p=7.24×10^−17^). There were also significant associations in different directions with numerous brain regional grey and white matter metrics, as well as psychological factors (Supplementary Table 7).

Full PheWAS outputs (regardless of the FDR threshold 0.05) for the ASD PRS and schizophrenia PRS are in Supplementary Tables 6 and 7. Handedness showed evidence for association with the ASD PRS, but this did not meet the FDR threshold 0.05 (multinomial logistic regression over three categories: right (n=28,703), left (n=3059), and ambidextrous (n=490), uncorrected p=0.001). This result was driven particularly by an association of higher ASD PRS with an increased rate of mixed-handedness (logistic regression beta=0.14, p=0.002), rather than left-handedness (beta=0.04, p=0.05), compared to right-handedness. Handedness was not associated with the schizophrenia PRS (p=0.63). ‘Volume of brain, grey+white matter’ was not associated with the ASD PRS (beta=−0.006, p=0.16), but showed evidence for a negative association with the schizophrenia PRS (beta=−0.02, uncorrected p=1.86×10^−5^).

As the schizophrenia PRS showed a negative association with brain size, we tested whether adjusting the 42 AIs for brain size by linear regression (in addition to all other covariates used in the primary analysis above) would alter the multivariate association of AIs with either PRS. However, both multivariate associations remained significant and largely unchanged (ASD PRS CCA r=0.03, p=2.25×10^−9^, schizophrenia PRS CCA r=0.04, p=2.57×10^−11^).

## Discussion

In this study of the UK Biobank brain imaging-genetic dataset, we found that PRS for ASD and schizophrenia were weakly correlated with each other, and each PRS was significantly associated with a distinct multivariate loading pattern across brain asymmetry measures. Contrary to expectations, neither PRS was associated with a more male-like or female-like pattern of brain asymmetry, after adjusting the AIs for sex. Thus, at least with respect to structural brain asymmetry, ASD polygenic risk does not appear to resemble increased masculinization, as might be expected according to the ‘extreme male brain’ theory of ASD^2^. Comparisons of cognitive profiles of ASD and schizophrenia to average sex differences may therefore be superficial, rather than reflecting an underlying risk dimension of etiological relevance. Furthermore, there was no evidence that multivariate patterns of brain asymmetry associated with the two PRS were anti-correlated, and therefore our results do not support concepts of ASD and schizophrenia as opposing disorders on a single neurobiological dimension (see Introduction). Rather, our findings support ASD and schizophrenia as largely, but not wholly, distinct disorders at the genetic and neurobiological levels.

The single asymmetry index that was noticeably concordant in its associations with both PRS was that of the pars opercularis surface area, a region of inferior frontal cortex that forms part of Broca’s classically defined language-production region^76^. Language is a left-lateralized function in most people^77^, and some leftward structural asymmetries of language-important regions, such as the pars opercularis, may reflect this functional laterality at the population level^78^. The pars opercularis showed reduced leftward asymmetry with higher PRS for both disorders, consistent with leftward asymmetry being the optimal structural organization. This region is likely to have contributed to the language-related terms that we observed for both PRS, in fMRI-based functional annotation of regions for which asymmetries were associated with the PRS. Therefore the two disorders may share an etiological link through altered structure and function of Broca’s region, consistent with impaired social communication in both disorders (see Introduction).

In addition to language, functional annotation indicated that brain regions for which asymmetries were associated with PRS for the two disorders were involved in executive functions. This was consistent with the phenome-wide association results, where a higher PRS for schizophrenia was most prominently associated with lower performance on tests requiring executive function (e.g. symbol-digit substitution and trail making). In contrast, the ASD PRS was significantly and positively associated with college or university qualifications, and also showed a positive trend of association with fluid intelligence (beta=0.04, uncorrected p=8.78×10^−4^), although the latter did not meet FDR correction at 0.05. A recent meta-analysis that compared cognitive performance across multiple domains in ASD and schizophrenia, in adulthood, also found the clearest differences in visuospatial perception, reasoning and problem solving, with ASD individuals performing better^79^. The ASD PRS was most significantly associated with difficulties following conversation in the presence of background noise, which may relate to a linguistic or social cognitive difficulty, rather than a sensory issue. Taken all together, the patterns of PRS-brain-behaviour associations in the UK Biobank suggest that largely distinct alterations of regional brain asymmetry may be functionally and etiologically relevant manifestations of high polygenic risks for ASD and schizophrenia, although future analyses in longitudinal data will likely be required to help clarify mediation and cause-effect relations.

The association effect sizes in this study were small, with canonical correlations no greater than 0.04, while the high degree of statistical significance arose through the large sample size. Macro-structural measures of regional asymmetry are relatively crude biological readouts, but it is possible that more substantial associations of disorder PRS will be detected with brain asymmetries measured at higher levels of resolution, for example in the abundances of specific cell or synaptic types, or gene expression levels^80^, with potentially more direct functional relevance. The brain asymmetry measures in this study all had heritabilities less than 10%, which means that any polygenic effects will necessarily be limited. In fact, most variance in brain asymmetries may be due to early developmental randomness^52,81–83^. The value of the present findings is not in terms of biomarkers for disorder risk, but rather in terms of biological clues into potentially relevant aspects of disorder etiology.

We found no evidence that brain size influenced the associations of either PRS with brain asymmetries. This may be expected as the AI (left−right)/((left+right)/2) is adjusted through its denominator for the bilateral magnitude of any given regional measure. We did observe that the schizophrenia PRS was associated with reduced brain size, consistent with average reductions in cortical surface area and thickness, and subcortical volumes, which have been reported to associate with schizophrenia by large-scale consortium analysis^84,85^. There was no association of the ASD PRS with brain size. This may reflect that the average age of the UK Biobank individuals in this study was 63 years. A previous study of 259 ASD individuals and 327 controls, aged up to 50 years, found relative brain overgrowth in ASD during infancy and childhood, followed by accelerated decline in size from adolescence onwards^86^. It is a limitation of the present study that the age of UK Biobank individuals is far older than the typical ages of onset of either ASD or schizophrenia. However, both disorders often persist throughout life^79^, and it is during adulthood that they can be most directly compared and contrasted in terms of affected brain traits and cognitive profiles.

The PRS for ASD and schizophrenia were weakly and positively correlated with each other in the present study (r=0.08), which is broadly consistent with a positive genetic correlation of roughly 0.2 between the two disorders^18^. Both PRS were also slightly higher in females than males. The UK Biobank is not fully representative of the population, with generally superior health and socioeconomic circumstances, and greater female participation^87^. Higher risk for psychiatric disorders has also been suggested to reduce participation in cohort studies^88^. Together, these observations suggest that the UK Biobank particularly under-represents males with high polygenic risks for ASD and schizophrenia. Regardless, when adjusting for sex, the multivariate associations of both PRS with brain asymmetries were highly significant, and not associated with either a more male-like or more female-like pattern.

We did not consider handedness as a confound variable in our analyses, because we were interested in associations of disorder PRS with structural brain asymmetries regardless of associations with other aspects of brain or behavioural asymmetry. Handedness may itself be a causally influenced variable by both disorder PRS and brain structural asymmetry, thus treating handedness as a confound variable could have induced collider bias into our analysis^89^. There was evidence that the ASD PRS was positively associated with non-right-handedness, consistent with an increased rate of this trait in ASD^42^.

In summary, we found that polygenic risks for ASD and schizophrenia have generally distinct associations with brain asymmetries. Findings were consistent with a particular involvement of regions important for language and executive functions, but there was no evidence that the two PRS were associated with overall opposing patterns of brain asymmetry, or with excessive masculinization or feminization of the brain. The most concordant association for the two PRS involved asymmetry of the pars opercularis, which supports the two disorders as sharing etiological features in the language domain. These findings contribute to an improved understanding of potentially shared and distinct etiological risks for ASD and schizophrenia.

## Acknowledgements

This research was funded by the Max Planck Society (Germany). The funders had no role in study design, data collection and analysis, decision to publish or preparation of the manuscript.The research was conducted using the UK Biobank resource under Application Number 16066, with Clyde Francks as the principal applicant. Our study made use of imaging-derived phenotypes generated by an image-processing pipeline developed and run on behalf of UK Biobank.

## Contributions

Z.S.: Conceptualization, methodology, analysis, visualization, original draft writing, review &editing. D.S.: Methodology, analysis, review &editing. C. F.: Conceptualization, direction, supervision, original draft writing, review &editing.

## Ethics declarations

The UK Biobank received ethical approval from the National Research Ethics Service Committee North West-Haydock (reference 11/NW/0382), and all of their procedures were performed in accordance with the World Medical Association guidelines. Informed consent was obtained for all participants.

The authors report no competing interests.

## Data Availability

The primary data used in this study are available via the UK Biobank website www.ukbiobank.ac.uk. Other publicly available data sources and applications are cited in the manuscript.

## Code availability

This study used openly available software and codes as cited in the Methods section.

